# Phylogenomic analyses of deep gastropod relationships reject Orthogastropoda

**DOI:** 10.1101/007039

**Authors:** Felipe Zapata, Nerida G. Wilson, Mark Howison, Sónia C.S. Andrade, Katharina M. Jörger, Michael Schrödl, Freya E. Goetz, Gonzalo Giribet, Casey W. Dunn

## Abstract

Gastropods are a highly diverse clade of molluscs that includes many familiar animals, such as limpets, snails, slugs, and sea slugs. It is one of the most abundant groups of animals in the sea and the only molluscan lineage that has successfully colonised land. Yet the relationships among and within its constituent clades have remained in flux for over a century of morphological, anatomical and molecular study. Here we re-evaluate gastropod phylogenetic relationships by collecting new transcriptome data for 40 species and analysing them in combination with publicly available genomes and transcriptomes. Our datasets include all five main gastropod clades: Patellogastropoda, Vetigastropoda, Neritimorpha, Caenogastropoda and Heterobranchia. We use two different methods to assign orthology, subsample each of these matrices into three increasingly dense subsets, and analyse all six of these supermatrices with two different models of molecular evolution. All twelve analyses yield the same unrooted network connecting the five major gastropod lineages. This reduces deep gastropod phylogeny to three alternative rooting hypotheses. These results reject the prevalent hypothesis of gastropod phylogeny, Orthogastropoda. Our dated tree is congruent with a possible end-Permian recovery of some gastropod clades, namely Caenogastropoda and some Heterobranchia subclades.

## 1. Introduction

Gastropoda, the clade of molluscs that includes snails, slugs, and their relatives, is hyperdiverse with respect to species number, morphology, habitat, and many other attributes. They radiated in marine, freshwater and terrestrial systems, and display extensive body plan disparity [1]. 32,000 to 40,000 marine species of gastropods have been described, but this is thought to only represent between 23 and 32% of the total estimated number of marine species of gastropods [2]. In addition, there is a large number of limno-terrestrial snails and slugs [3], many of which are threatened to a degree unparalleled among other invertebrate groups [4]. The overall magnitude of the gastropod diversity is extremely hard to estimate; in a survey of a New Caledonian coral reef lagoon, gastropods represented almost 80% of the 2,738 species of molluscs found (excluding cephalopods) [5], with many undescribed species.

Gastropods are characterised by having a single shell and an operculum, at least in the larval stage, and by undergoing torsion during development. They range in size from less than 1 mm to almost 1 m, and their shell has been modified enormously in many groups, including the common coiled and torted (usually dextrally) snail-like, the highly convergent limpet-like, or the rare tubular or even bivalved shells [6]. Many lineages have reduced the shell or it has been entirely lost.

Gastropod relationships have been at the centre of molluscan research and have been in flux for decades (see Fig. 1). Many authors have employed cladistic methods to analyse morphological data [6–10]. This work supports the monophyly of gastropods and the division of the group into five main clades, Patellogastropoda, Vetigastropoda, Neritimorpha, Caenogastropoda and Heterobranchia, in addition to the less-understood Cocculinida and the so-called ‘hot-vent taxa’ (Peltospiridae and Cyathermiidae). The first numerical cladistic analysis included 117 morphological characters coded for 40 taxa, dividing gastropods into Eogastropoda (Patellogastropoda and Neolepetopsoidea; but several authors now find Neolepetopsoidea nested within Patellogastropoda [11]) and Orthogastropoda (all remaining gastropods) (Fig. 1a: [10]). Other well-supported clades recovered in these analyses included Patellogastropoda, Vetigastropoda, Neritimorpha, Caenogastropoda and Heterobranchia, the latter two forming the clade Apogastropoda (Fig. 1a-f). However, the Eogastropoda/Orthogastropoda division has not been supported in other analyses combining morphology with molecules (Fig. 1b: [6]), or in molecular analyses (e.g., [12–18]), which tend to find support for Thiele’s [19] clade Archaeogastropoda (with or without Neritimorpha) (see Fig. 1 for a summary of hypotheses).

**Figure 1.**
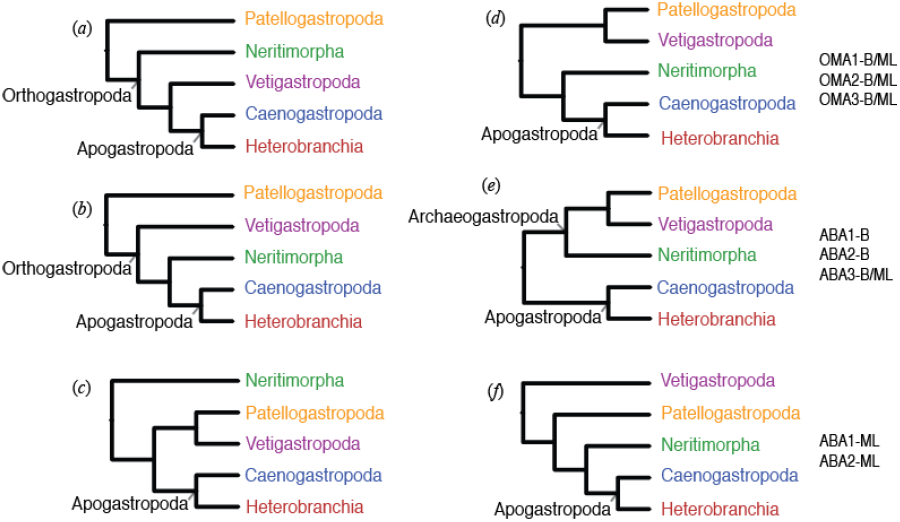
Hypotheses for the internal relationships of Gastropoda. Not all listed studies find monophyly of all taxa, as Vetigastropoda is often paraphyletic or diphyletic in earlier studies. Apogastropoda (= Caenogastropoda + Heterobranchia) is monophyletic in nearly all published studies. Hypotheses on the left do not have support from the analyses presented here. Hypotheses on the right are consistent with the analyses presented here, and differ only in their rooting. Matrix construction (ABA, OMA), subsampling strategy (1,2,3), and inference method (Bayes, ML) supporting each of these hypotheses is indicated.

Heterobranchia comprises the most diverse and ecologically widespread gastropod clades including the informal groups allogastropods, opisthobranchs and pulmonates [20]. With conservative estimates suggesting more than 40,000 species, heterobranchs are abundant in habitats ranging from the benthic realm to pelagic, intertidal to deep sea, tropical to polar, and freshwater to terrestrial [3, 21]. These transitions are not evenly spread across lineages, and the concomitant morphological specialisations have made defining homologies difficult in many cases [e.g., 22]. Although a consensus of relationships among heterobranch groups is emerging [23, 24], and Panpulmonata [see 25] has been recently supported [26], the monophyly and relationships of other higher taxa (e.g., Nudipleura, Tectipleura) have not been evaluated with next generation data.

In this study, we address the evolution of Gastropoda and evaluate the relationships among and within major clades in this group by creating a comprehensive taxonomic dataset from 40 novel transcriptomes and 16 publicly available genomes or transcriptomes. Using information from multiple nuclear-protein coding genes provide large amounts of data that can provide key phylogenetic insight [e.g., 26] as well as facilitate several aspects of phylogenetic inference.

## 2. Materials and Methods

### (a) Taxon sampling, RNA isolation, and Sequencing

We collected new transcriptome data for 40 species, including 34 gastropods and 6 other molluscs. All new datasets are paired end Illumina reads, except for single end Illumina datasets for *Hinea brasiliana*, *Philine angasi*, and *Strubellia wawrai*. Samples were prepared for sequencing with TruSeq RNA Sample Prep Kit (Illumina) or a previously described custom protocol [27]. We deposited all these new sequence data, along with associated specimen collection information, voucher accession numbers, RNA extraction methods, and library preparation details, in NCBI Sequence Read Archive (BioProject PRJNA253054). Vouchers for most specimens were deposited at the Museum of Comparative Zoology, Harvard University (Cambridge, Massachusetts, USA) and Scripps Institution of Oceanography (La Jolla, California, USA). The publicly available data for *Siphonaria pectinata* is here shown in the figures as *Siphonaria naufragum*, according to a recent revision [28].

### (b) Data analyses

These data were analysed in combination with publicly available data for 16 additional species to generate 56-taxon matrices. All Illumina reads (new and publicly available) were assembled with Agalma (versions 0.3.4-0.3.5) [29], 454 datasets were assembled externally with Newbler (versions 2.3 and 2.5p1), and gene predictions from *Lottia gigantea* [30] and *Pinctada fucata* [31] were imported directly into Agalma. Source code for most analysis steps as well as sequence alignments, tree sets, summary trees, and voucher information are available in a git repository at https://bitbucket.org/caseywdunn/gastropoda.

Two methods were used to generate the supermatrices within Agalma. In method 1, after assembly, translation and removal of mtDNA loci, the sequences from all taxa were compared to each other using an All-By-All BLASTP search, and a phylogenetic approach to identify orthologous sequences (see [32]). We refer to this method as ABA. In method 2, the sequences from all taxa were compared using OMA v0.99t [33] to directly assign sequences to groups of orthologs using an entirely phenetic approach (see [34]). We refer to this method as OMA.

For each method (ABA and OMA) we constructed three progressively smaller and denser amino acid supermatrices, creating a total of six matrices (Fig. 2). Supermatrix 1 was constructed by concatenating all ortholog sequences until the cumulative gene occupancy was 50% (49,752 sites/862 loci for ABA and 190,752 sites/1,245 loci for OMA; 425 loci in common between ABA and OMA) and then removing *Pyropelta* sp. and *Paralepetopsis* sp., which were poorly sampled. Supermatrix 2 was constructed by removing taxa with less than 20% gene occupancy from Supermatrix 1. The removed taxa include *Haliotis kamtschatkana*, *Perotrochus lucaya*, *Littorina littorea*, *Siphonaria naufragum*, *Chaetoderma* sp., and *Pomacea diffusa* for both OMA and ABA matrices, as well as *Amphiplica gordensis* for the ABA Supermatrix 2. This taxon was removed from the ABA Supermatrix 2 bootstrap replicates, and the ABA Supermatrix 2 posterior probability tree sets prior to summary so that they could be consistently displayed (Supplementary Figs. 1-2). Supermatrix 3 was constructed by trimming genes from Supermatrix 2 until the cumulative gene occupancy reached 70% (15,735 sites/300 loci for ABA and 45,084 sites/364 loci for OMA; 110 loci in common between ABA and OMA).

**Figure 2.**
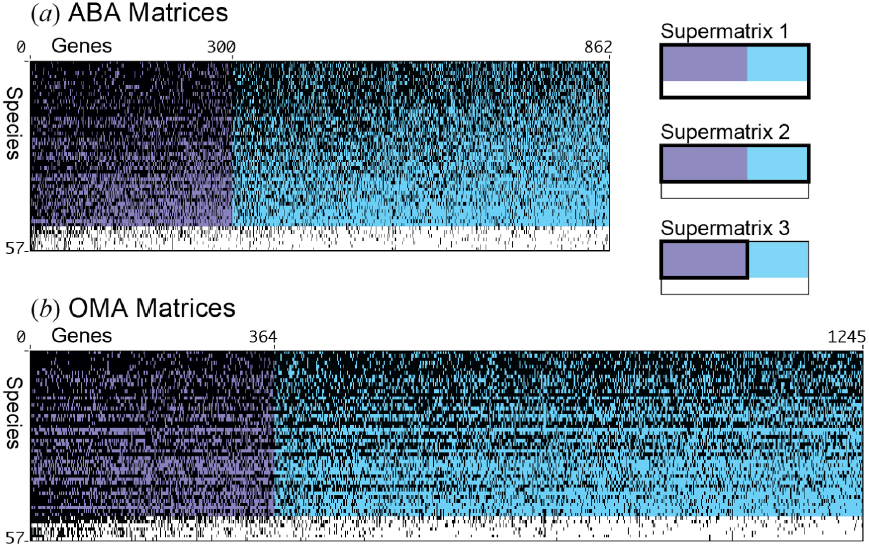
The six matrices that were considered here. Supermatrices were assembled with two methods, ABA (a) and OMA (b). Three matrices were constructed for each of these methods. Supermatrix 1 is the full set of genes and species. From Supermatrix 1, Supermatrix 2 is constructed as a subset of the best sampled species. From Supermatrix 2, Supermatrix 3 is constructed as a subset of the best sampled genes. See methods for additional details. Black indicates sampled genes for each taxon. Genes and species are sorted by sampling, with the best sampled in the upper left.

We inferred phylogenetic relationships using both maximum likelihood (ML) and Bayesian approaches, for a total of 12 phylogenetic analyses on the six supermatrices. For ML, we used ExaML v1.0.11 [35] with a WAG+Γ model of amino acid evolution. Bootstrap values were estimated with 200 replicates. Bayesian analyses were conducted with PhyloBayes MPI v1.4e [36] using the CAT-Poisson model of amino acid evolution. Two independent MCMC chains were run on each matrix adjusting the number of cycles until convergence was achieved. Convergence was determined with time-series plots of the likelihood scores, time-series plots of the cumulative split frequencies, maximum bipartition discrepancies across chains < 0.1, and an estimated effective sample size of tree likelihoods of at least 100. Post burn-in sampled trees were combined and summarised with a majority rule consensus tree.

Tree dating was conducted with MCMCTree v4.7 [37] using the approximate likelihood calculation algorithm [38], and the WAG+Γ model of evolution. A birth-death speciation process was specified as tree prior with default parameters (death and growth rate parameters equal 1, and sampling parameter equals 0). Rate heterogeneity among lineages was modelled using an uncorrelated lognormal relaxed molecular clock [39] with a diffuse gamma Γ(1,1) prior for the substitution rate and the rate-drift parameter. We used fossil calibrations to set prior densities on the ages of five nodes (Supp. Fig. 3) using minimum soft bounds with a left tail probability of 2.5% [40]. Because MCMCTree always needs a calibration point on the root [37], we used 550 my (ca. Terreneuvian; see [41]) to set a prior density on the root age using a maximum soft bound with 2.5% tail probability. We ran MCMCTree twice each time for 1.2e + 07 generations, sampling every 1.0e + 03, and discarding 20% of the samples as burn-in. Convergence was determined with time-series plots of the likelihood scores and assessing for correlation of divergence times between runs.

### (c) Hypothesis testing for Orthogastropoda

We statistically compared the Orthogastropoda hypothesis to our maximum likelihood tree using the SOWH test [42]. To carry out this analysis, we used SOWHAT [43] specifying a constraint tree and the WAG+Γ model on supermatrix 1 (OMA). We used the automatic stopping criterion implemented in SOWHAT to determine an appropriate sample size for the null distribution.

## 3. Results and Discussion

### (a) Deep relationships among major gastropod clades

Our data sets strongly support the monophyly of gastropods. This result is not surprising in itself, but has only recently been supported by molecular analyses of large data sets [32, 44, 45; see also 14, 15, 17, 18], or in the total evidence analysis of Aktipis *et al.* [6] Our analyses also support the monophyly of all major gastropod clades represented by multiple taxa: Vetigastropoda, Neritimorpha, Caenogastropoda and Heterobranchia (Fig. 3a). Patellogastropoda is represented by a single species, so its monophyly could not be evaluated. The deep internal relationships of gastropods therefore can be reduced to a 5-taxon problem (Fig. 1, 3b). Our 12 analyses (two inferences methods on two types of supermatrices each subsampled in three different ways) all recover the same unrooted ingroup relationships for these five clades (Fig. 3b, Supplementary Figs. 1b, 2b). These ingroup relationships are strongly supported by all methods except the ABA ML analyses, which have lower support than the other methods for a bipartition Vetigastropoda + Patellogastropoda (58%, 75%, and 56% for Supermatrices 1, 2, and 3) and recover Vetigastropoda + Neritimorpha in a minority of replicates. The lower support in these analyses may be due to the poor sampling of Patellogastropoda. These ingroup relationships allow us to reject the hypotheses for gastropod relationships indicated in Fig. 1a and 1b.

**Figure 3.**
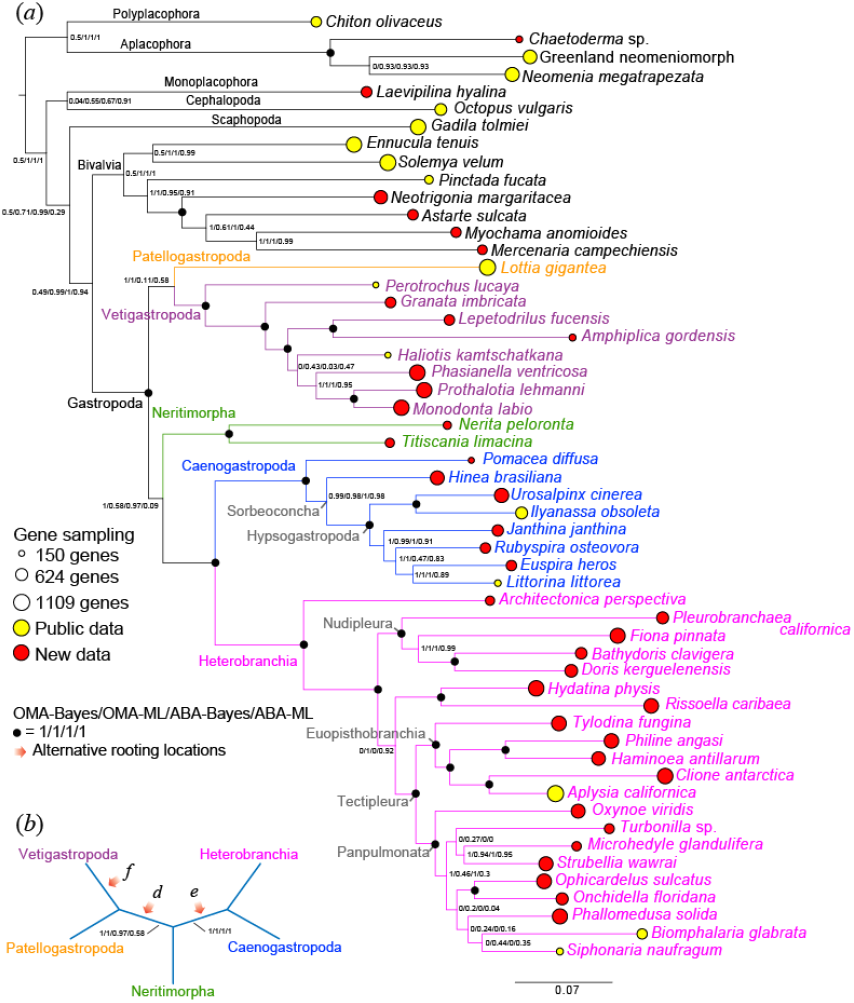
Summary tree for analyses of Supermatrices 1. (a) Rooted phylogram of the maximum likelihood OMA analysis, including outgroup taxa. Branch support values are shown on descendent nodes. The areas of the lollipops, which are centered on the branch tips, are proportional to the number of genes sampled in OMA supermatrix 1. (b) Unrooted cladogram of the ingroup taxa. Branch support values are shown, and alternative rooting locations are indicated with orange arrows. These support values were calculated by removing the outgroup taxa from the tree sets used to generate (a) and regenerating consensus trees. The letter on the rooting arrow corresponds to the hypotheses shown in Fig. 1.

Although the ingroup relationships found broad consistent support, the rooting of gastropods is still not well resolved. Our results are congruent with three possible rootings (orange arrows in Fig. 3b, Supplementary Figs. 1b, 2b). This is akin to other recalcitrant animal phylogeny questions, including the root of Metazoa [e.g., 46, 47] and the root of arthropods [48]. Though the hypothesis indicated in Fig. 1c is compatible with the ingroup relationships supported here, we never recover this rooting and it can be excluded. This reduces the possible alternatives for deep gastropod relationships to the three hypotheses (Fig. 1d-f). Two of these remaining hypotheses have been proposed before [6, 14]; the other (Fig. 1f) is recovered for the first time here.

The rejection of several widely held hypotheses for deep gastropod phylogeny (Figs. 1a-c) has major implications for the understanding of gastropod evolution. All our analyses reject the Orthogastropoda hypothesis (a clade comprised of Vetigastropoda, Neritimorpha, Caenogastropoda and Heterobranchia) and the placement of Patellogastropoda as the sister group to other gastropods (Figs. 1a, 1b). Even in the minority of ABA ML replicates that recover an ingroup partition Vetigastropoda + Neritimorpha, the rooting is inconsistent with Orthogastropoda. The broadly accepted Orthogastropoda hypothesis has been proposed in multiple configurations [e.g., 6, 9, 10, 49, 50]. The placement of Patellogastropoda as the sister group to Orthogastropoda has been driven by considerable anatomical research. One potential character supporting this placement is the ciliary ultrastructure of the cephalic tentacles, which also occurs in Bivalvia and Solenogastres but is lacking from other gastropods [51]. In this scenario, this character is plesiomorphic for Mollusca, retained in Patellogastropoda and was lost a single time in Orthogastropoda. However, because enforcing the monophyly of Orthogastropoda is significantly worse (SOWH test: N = 152, Δ-likelihood = 374.0137, *p* = 0) than our most likely tree (Fig. 3), our results indicate that this character may be convergent between Patellogastropoda and outgroup taxa, or was lost more than once within Gastropoda. We also reject another recent hypothesis for gastropod rooting, the sister group relationship of Neritimorpha to other Gastropoda (Fig. 1c, [e.g., 15, 17, 18]).

Our reduction of deep gastropod phylogeny to three alternative hypotheses (Figs. 1d-f) clarifies multiple open questions. All three of these hypotheses include the monophyly of Apogastropoda (= Heterobranchia + Caenogastropoda), reinforcing this widely accepted aspect of gastropod relationships. Other relationships supported here have been found earlier (Fig. 1d-e) (see review by Aktipis *et al*., [6]; for different views see e.g., the gastropod classification by Bouchet & Rocroi [52] and the mitogenomic study by Grande *et al.* [6, 14, 53]). However, it is now acknowledged that mitogenomic data are not appropriate for resolving deep gastropod relationships [54]. To our knowledge, no molecular analysis supported the placement of Vetigastropoda as sister group to all other gastropods.

Because we ran twelve phylogenetic analyses, we can explore the differences in support between these three alternative hypotheses for gastropod rooting across inference method (Bayes and ML), matrix construction method (OMA and ABA), and matrix subsampling (Supermatrices 1, 2, and 3; Fig. 2). Matrix subsampling had little effect on deep relationships. Analyses of Supermatrix 1 (Fig. 3) and Supermatrix 2 (Supplementary Fig. 1) were consistent with all three rooting positions (Fig. 1d-f). Analyses of Supermatrix 3 (Supplementary Fig. 2) found support for only two of these rootings (Fig. 1d,f). Unlike the other analyses, it did not recover Apogastropoda as the sister group to all other gastropods (Fig. 1e). This particular hypothesis (Fig. 1e) is interesting because it includes Archaeogastropoda, which was proposed nearly a century ago by Thiele [19]. Bayesian analyses recovered Neritimorpha as the sister group to Apogastropoda (Fig. 1d,f) in all analyses, but ML analyses found very low support for this relationship. Analyses of OMA matrices provided strong support for a clade comprised of Patellogastropoda and Vetigastropoda (Fig 1d,e), but analyses of ABA matrices do not.

These results suggest clear strategies for distinguishing between the remaining hypotheses for deep gastropod relationships. Since these hypotheses differ only in their rooting, improved outgroup sampling will be critical. To maximize gene sampling and matrix density, we limited our sampling of non-gastropod molluscs to those for which Illumina data [32] or genomes are available. Future analyses of gastropod relationships will need to include more outgroups to resolve the remaining open questions. Previous phylogenomic analyses of molluscs that also included extensive 454 and Sanger data [32, 44] had much broader non-gastropod sampling but minimal gastropod sampling. The rooting of gastropods was not fully supported in these analyses either, but the strongly supported ingroup relationships are compatible with the three hypotheses supported here. In addition, improved sampling of Patellogastropoda (here represented by a single species with a complete genome) and Neritimorpha, and the addition of the unsampled Neomphalina and Cocculiniformia will be critical.

### (b) Relationships within major gastropod clades

Though our sampling is focused on resolving deep relationships between the major gastropod clades, our results do find strong support for some previously unresolved relationships within Vetigastropoda, Caenogastropoda, and Heterobranchia.

A key question within Vetigastropoda is the placement of Pleurotomarioidea (*Perotrochus* in our analyses), which appears outside Vetigastropoda in some previous studies [14, 15]. Here we find strong support for the placement of Pleurotomarioidea as the sister group to all other vetigastropods (Fig. 3a), resolving this issue. We also resolve the position of Seguenzioidea (*Granata*) as the next clade to diverge from other vetigastropods (Fig. 3a, Supplementary Figs. 1-2). Our analyses also recover a well supported clade of deep-sea taxa (Pseudococculinidae [*Amphiplica*] and Lepetodriloidea [*Lepetodrilus*]; Fig. 3a). We also find strong support (Fig. 3a, Supplementary Figs. 1-2) for a clade comprised of Phasianellidae (*Phasianella*) and Trochoidea (*Prothalotia* and *Monodonta*). The position of *Haliotis* is not resolved (Fig. 3a), perhaps due to relatively poor gene sampling.

Caenogastropoda is a megadiverse clade comprising about 60% of living gastropod species [55], so our limited sampling can address only a small fraction of open questions about internal relationships of this group. The relationships we can test are largely in agreement with prior morphological [e.g., 10, 55–57] and molecular [e.g., 58–60] analyses. We find a sister group relationship of Ampullarioidea (represented by *Pomacea*) to Sorbeoconcha, which comprise the remaining sampled caenogastropods (Fig. 3a). Within Sorbeoconcha, Cerithioidea (*Hinea*) is the sister group to Hypsogastropoda, the latter dividing into a siphonate (in our case the two Neogastropoda: *Urosalpinx* and *Ilyanassa*) and an asiphonate group (*Janthina*, *Littorina*, *Euspira* plus *Rubyspira*), similar to Ponder *et al.* [55].

The basic structure of internal Heterobranchia relationships has only recently gained some agreement [23, 24]. Our strong support for the placement of *Architectonica* as sister group to the other sampled heterobranchs is consistent with most other analyses [see 24]. Nudibranchia (*Fiona* + *Bathydoris* + *Doris*) and Nudipleura (*Pleurobranchaea* + Nudibranchia) were monophyletic [61, 62], despite some suggestion that Pleurobranchoidea may not be the sister group to the remaining nudibranch lineages [26]. Our results recovered a monophyletic Cephalaspidea (*Philine* + *Haminoea*), sister group to Anaspidea (*Aplysia*) + Pteropoda (*Clione*). Umbraculoidea (*Tylodina*) was sister group to Cephalaspidea + Anaspidea + Pteropoda; all four taxa together represent the well supported Euopisthobranchia (Fig. 3a, Supplementary Figs. 1-2). We find support for Panpulmonata (Fig. 3a, Supplementary Figs. 1-2), but their internal relationships are mostly unresolved and clearly require future attention. Like previous Sanger sequencing-based studies, our analyses consistently recover a Panpulmonata + Euopisthobranchia clade, or Tectipleura [25, 63]. The relationship of Tectipleura to other heterobranchs has been of particular interest. We recover two conflicting hypotheses for these relationships, neither of which has been previously proposed. Our likelihood analyses place the unnamed clade Rissoelloidea (*Rissoella*) + Acteonoidea (*Hydatina*) as the sister group to Tectipleura (Fig. 3a, Supplementary Figs. 1-2). Our Bayesian analyses, however, place Nudipleura and this Rissoelloidea + Acteonoidea clade together with strong support, and place this clade as sister to Tectipleura (Supplementary Fig. 3). Previous analyses have instead favored Euthyneura, a clade comprised of Tectipleura and Nudipleura (but excluding Rissoelloidea + Acteonoidea). We do not recover Euthyneura in any of our analyses. Tectipleura is united by a monaulic reproductive system, but even Euthyneura is not entirely defined by euthyneury, as reversals are known in several subgroups (e.g., Rhodopemorpha [64, 65]). Rissoelloidea + Acteonoidea and Euthyneura share giant neurons in macroscopic animals [64], and if necessary, a simple redefinition of the taxon Euthyneura to include Rissoelloidea + Acteonoidea would maintain stability.

### (c) Chronogram

Our dated phylogeny (Fig. 4) shows a Cambrian origin of stem gastropods with crown diversification into its five main lineages during the Ordovician to the Devonian, as well shown in the gastropod fossil record [66]. From the well-sampled groups, crown Vetigastropoda diversified first, around the Devonian-Carboniferous, followed by Neritimorpha and Heterobranchia at similar periods. Crown Caenogastropoda seem to have diversified later, around the Permian-Triassic, perhaps initiating its explosive diversification after the end-Permian mass extinction ca. 254 Ma, responsible for the extinction of 95–99% of marine species and to change the ecosystems and their faunal composition forever. Such drastic post extinction diversifications have been recently shown for other modern clades of marine organisms (e.g., Crinoidea [67]; Protobranchia [68]). This could also explain other explosive radiations in gastropods, especially within the euthyneuran Heterobranchia clades such as Nudipleura (Fig. 4). Denser sampling will however be required to derive accurate diversification curves to test these hypotheses.

**Figure 4.**
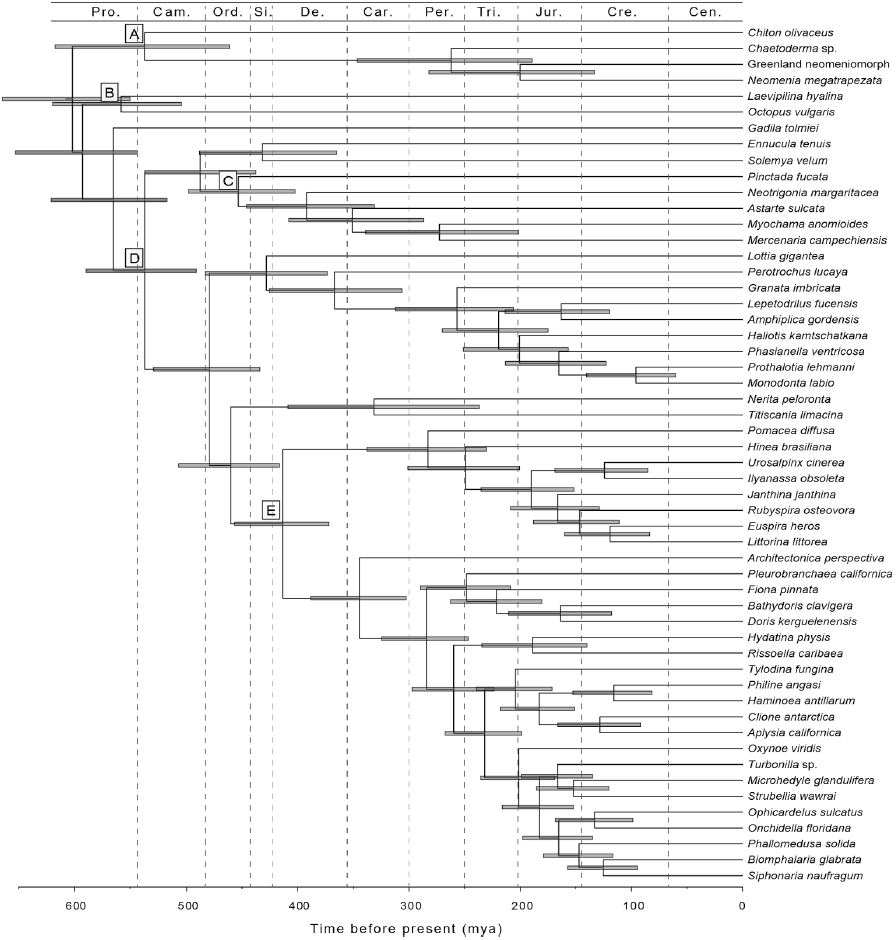
Chronogram with estimates of divergence times for internal nodes. Bars correspond to 95% credibility intervals. Fossil constraints were set on nodes A-E: node A, 231 my (*Leptochiton davolii* [68]); node B, 505 my (*Plectronoceras cambria* [69]); node C, 475 my (*Glyptarca serrata*, Arenigian [70]); node D, 530 my (*Fordilla troyensis* from the Tommotian of Siberia [71, 72, 73]); node D, 418 my (Sublitoidea [74]). We used 550 my (ca. Terreneuvian; see [41]) to set a prior density on the root age using a maximum soft bound with 2.5% tail probability. Geological periods abbreviated on top: Pro.: Proterozoic, Cam.: Cambrian, Ord.: Ordovician, Si.: Silurian, Dev.: Devonian, Car.: Carboniferous, Per.: Permian, Tri.: Triassic, Jur. Jurassic, Cre. Cretaceous, Cen.: Cenozoic.

## Acknowledgments

This research was supported by the US National Science Foundation through the Systematics Program (awards 0844596, 0844881 and 0844652) and the Alan T. Waterman Award. Field work was supported by the US National Science Foundation through the Assembling the Tree of Life program BivAToL grant (award 0732903), and by the German Research Foundation (SCHR667/9-1 and 13-1). Sequencing at the Brown Genomics Core facility was supported in part by NIH P30RR031153 and NSF EPSCoR EPS-1004057 and sequencing at the Harvard Center for Systems Biology was supported with internal funds from the Museum of Comparative Zoology, and by the Volkswagen Foundation to KMJ. Data transfer was supported by NSF RII-C2 EPS-1005789. Analyses were conducted with computational resources and services at the Center for Computation and Visualization at Brown University, supported in part by the NSF EPSCoR EPS-1004057 and the State of Rhode Island. Thanks to Alicia R. Pérez-Porro, Vanessa González and Ana Riesgo for laboratory assistance, Stephen Smith for preliminary data analyses, Samuel Church for help running SOWHAT, and Robert Vrijenhoek at Monterey Bay Aquarium Research Institute (MBARI) for providing the *Rubyspira* and *Amphiplica* samples.

## Author contributions

NGW, GG, and CWD conceived of and designed the study. NGW, GG, KJ, MS, and FEG collected samples. SCSA, FEG, KJ and NGW prepared samples for sequencing. FZ designed and ran analyses. MH implemented software and assisted with data management. GG, FZ, NGW, and CWD wrote the manuscript. All authors discussed / contributed to the final manuscript version.

## Data accessibility

- Raw sequence data: NCBI Sequence Read Archive BioProject PRJNA253054, accesion numbers: SRR1505101-SRR1505105, SRR1505107-SRR1505141.
- Analysis scripts, phylogenetic alignmets, tree sets, summary trees, and voucher information: https://bitbucket.org/caseywdunn/gastropoda. The most recent commit at the time of submission is available at: https://bitbucket.org/caseywdunn/gastropoda/src/b93fce3bf8e90cc0124327f5f7d3d0353ee4d295
- Phylogenetic data also available at: http://dx.doi.org/10.5061/dryad.5bc98

## Supplementary Files

**Supplementary Figure 1.**
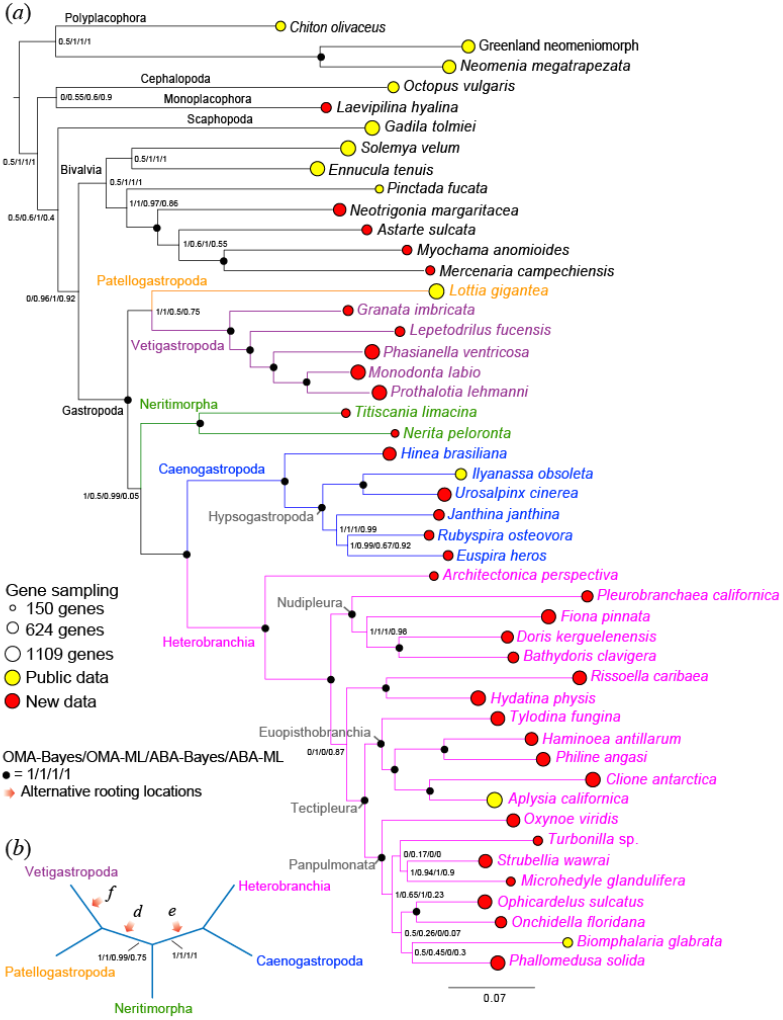
Summary tree for analyses of Supermatrices 2. See Figure 3 legend for description of layout.

**Supplementary Figure 2.**
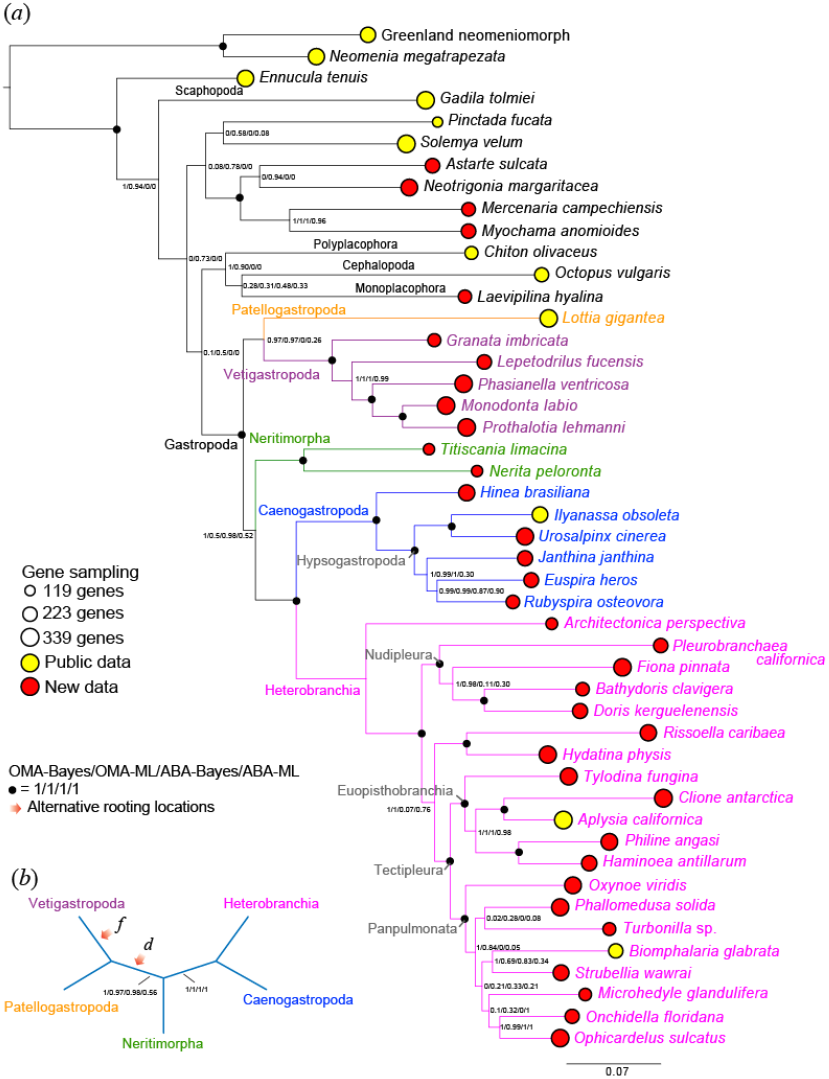
Summary tree for analyses of Supermatrices 3. See Figure 3 legend for description of layout.

**Supplementary Figure 3.**
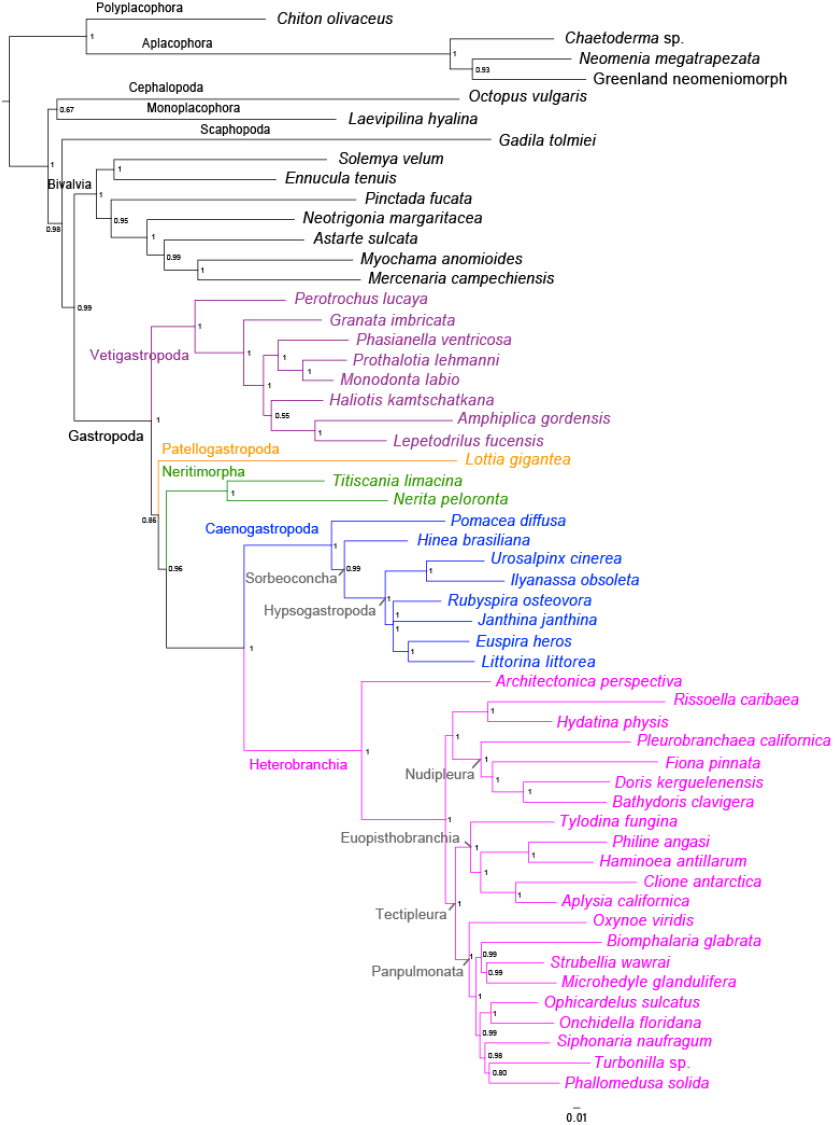
Phylogram of the Bayesian analysis of ABA Supermatrix 1. Tree sets for other analyses are available at DataDryad.

